# Paternal folate deficiency reveals meiosis as a metabolic sensing window in the male germline

**DOI:** 10.64898/2026.04.22.720272

**Authors:** Jasmine M. Esparza, Rohan V. Kumar, Mengwen Hu, Yasuhisa Munakata, Saanvi Jain, Akihiko Sakashita, So Maezawa, Richard M. Schultz, Satoshi H. Namekawa

## Abstract

Folate-dependent one-carbon metabolism supplies methyl donors required for chromatin modification, yet how metabolic conditions shape epigenome establishment in the male germline remains poorly understood. Here, using a folate-deficient mouse model, we identify meiotic prophase I as a metabolically sensitive window in the male germline. By integrating in vivo germline analysis with bulk and single-cell transcriptomic and epigenomic profiling, we show that folate deficiency disturbs transcriptional programs in pachytene spermatocytes and preferentially perturbs CpG island (CGI)-associated promoters, which are characterized by bivalent H3K4me3 and H3K27me3 during meiosis. Consistent with this selective vulnerability, active chromatin marks, including H3K4me3 and H3K27ac, are markedly reduced at CGI-associated promoters under folate-deficient conditions. Notably, loci that later exhibit altered H3K4me3 enrichment in mature sperm show earlier chromatin perturbations during meiosis, suggesting that these sperm epigenomic alterations may originate during meiotic development. Together, these findings establish a mechanistic link between paternal folate deficiency and dynamic epigenomic remodeling of CGI-associated chromatin in the male germline.

## Introduction

The importance of paternal health in shaping offspring outcomes has gained increasing attention in recent years (Rando & Simmons, 2015; Rotimi & Singh, 2024). In humans, poor diet, lifestyle factors, and environmental exposures can negatively affect sperm parameters such as sperm quality and fertility (Eisenberg *et al*, 2014; Ferramosca & Zara, 2022; Gaskins *et al*, 2012; Giahi *et al*, 2016; Rotimi & Singh, 2024). Beyond their role as carriers of genetic information, sperm convey epigenetic information linked to gene expression during early embryonic development and can influence offspring health (Kimmins & Sassone-Corsi, 2005; Lambrot *et al*, 2013; Lismer *et al*, 2021; Siklenka *et al*, 2015). Nutritional status can shape the epigenome by modulating the availability of metabolites required for chromatin-modifying reactions (Van den Veyver, 2002), providing a mechanistic link between paternal diet and germline epigenetic regulation. However, the developmental origins of diet-induced epigenomic information carried by sperm remain a critical gap in our understanding. Although germline epigenetic reprogramming during fetal development is a well-established window of vulnerability (Hayashi *et al*, 2012; Hill *et al*, 2018; Seisenberger *et al*, 2012), accumulating evidence indicates that environmental exposures and lifestyle factors in adult males also shape sperm epigenetic states and offspring phenotypes (Carone *et al*, 2010; Lambrot *et al*., 2013; Siklenka *et al*., 2015; Watkins *et al*, 2018).

Spermatogenesis entails extensive chromatin remodeling and epigenomic reprogramming to establish germ cell-specific transcriptomes required for male fertility (Alavattam *et al*, 2024; Hasegawa *et al*, 2015; Kitamura *et al*, 2025a; Kitamura & Namekawa, 2024, 2025; Kitamura *et al*, 2025b; Lin *et al*, 2025; Maezawa *et al*, 2020b; Maezawa *et al*, 2018b; Maezawa *et al*, 2021; Rabbani *et al*, 2022; Sin *et al*, 2015a). These processes are metabolically demanding and require tight coordination of growth, replication, differentiation, and chromatin regulation. Notably, meiotic prophase progression imposes particularly high metabolic and chromatin regulatory demands, driven in part by robust transcriptional activity in pachytene spermatocytes (Bolcun-Filas *et al*, 2011; Maezawa *et al*, 2020a; Soumillon *et al*, 2013), suggesting that meiotic prophase represents a critical window for sensing nutritional perturbations. Nevertheless, whether meiotic prophase functions as a metabolic checkpoint that translates nutritional status into stable germline epigenomic states remains unclear.

One-carbon metabolism, supported by dietary folate (vitamin B9) intake, supplies methyl donors such as S-adenosylmethionine (SAM) for DNA and histone methylation reactions underlying epigenomic regulation, thereby linking nutritional status to chromatin regulation (Petrova *et al*, 2023). Folate, obtained from the diet as naturally occurring folates or as synthetic folic acid from fortified foods and supplements, must be acquired exogenously (Berry *et al*, 2010; Crider *et al*, 2022). Perturbations in folate availability may therefore represent a physiologically relevant challenge to the fidelity of germline epigenetic programming during spermatogenesis. Although the benefits of maternal folate supplementation are well established (Jankovic-Karasoulos *et al*, 2021; Liu *et al*, 2020; van Uitert & Steegers-Theunissen, 2013; Yang *et al*, 2022), the role of paternal folate status in shaping germ cell competence and offspring health has received comparatively less attention, despite evidence linking paternal folate deficiency to adverse reproductive and developmental outcomes (Hoek *et al*, 2020; Lambrot *et al*., 2013; Lismer *et al*., 2021; Ly *et al*, 2017). Paternal folate deficiency alters histone H3 lysine 4 trimethylation (H3K4me3) patterns in mature sperm and is associated with offspring phenotypes (Lismer *et al*., 2021). Whether these alterations arise during late sperm maturation or earlier in spermatogenesis remains unknown.

Here, we integrate histological, bulk and single-cell transcriptomic, and epigenomic analysis to define the developmental timing and chromatin contexts in which paternal folate deficiency perturbs spermatogenesis. We identify meiotic pachytene spermatocytes as a particularly vulnerable stage in which folate deficiency disrupts epigenomic regulation, preceding aberrant epigenetic states in mature sperm. Together, these findings establish meiotic prophase as a metabolic sensing window in the male germline and link paternal nutrition to stage-specific germline epigenomic reprogramming.

## Results

### Folate deficiency impairs the germ cell population beginning at meiosis

To examine how paternal nutrition influences stage-specific epigenetic regulation during spermatogenesis, we used a paternal folate–deficient (FD) mouse model previously established for folate-induced epigenetic inheritance (Lismer *et al*., 2021) (Fig. 1A). Breeders and weanlings were maintained on a folate–sufficient (FS) diet until male littermates were weaned at 3 weeks-of-age and then randomly assigned to FS or FD diets until 14 weeks-of-age (Fig. 1B). In mice, differentiation of spermatogonia into mature sperm requires ∼35 days (Lismer *et al*., 2021) (Fig. 1C, D). Thus, 11 weeks of dietary exposure ensured that multiple complete cycles of spermatogenesis—from undifferentiated spermatogonia to spermatozoa—occurred under folate dietary stress. A previous study demonstrated that an FD diet reduced intratesticular folate levels (Lismer *et al*., 2021), confirming that germ cells experience folate limitation during spermatogenesis. Consistent with a prior report (Lambrot *et al*., 2013), testis-to-body weight ratios were comparable between FD and FS mice, although body weight—but not testis weight—was slightly increased in FD mice (Fig. EV1A–C). Weekly food consumption per mouse was comparable between FD and FS groups, indicating that differences in body weight were not explained by differences in food intake (Fig. EV1D). Notably, several human studies have also reported associations between folate deficiency and increased body mass index or body weight (Jafari *et al*, 2023; Mlodzik-Czyzewska *et al*, 2020; Pereira *et al*, 2019).

**Figure 1.**
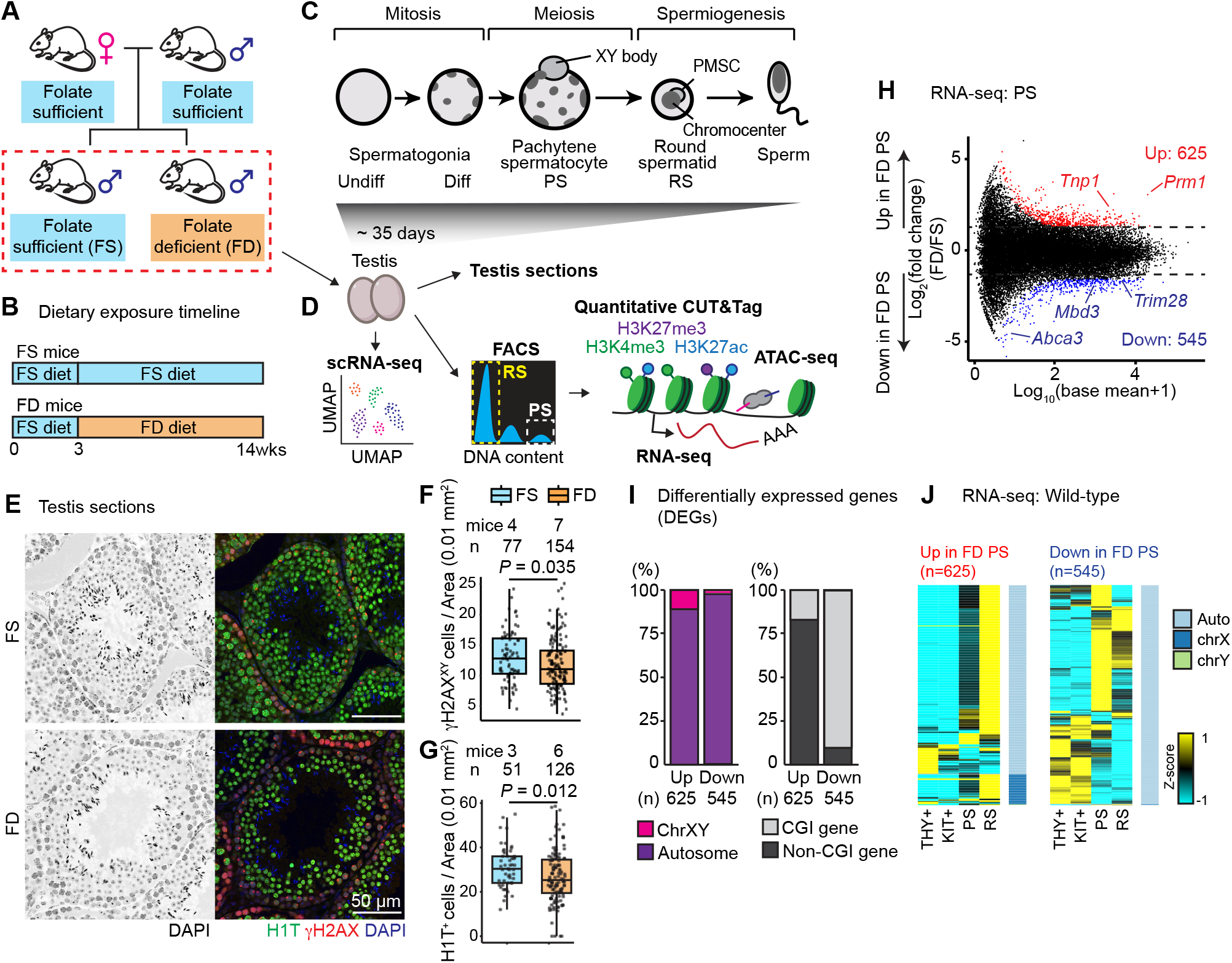
Folate deficiency affects spermatogenesis and disturbs transcriptional programs in pachytene spermatocytes. (A) Schematic of the folate-deficient (FD) dietary model. Breeder mice were maintained on a folate-sufficient (FS) diet. (B) Dietary exposure timeline. Littermate males were maintained on an FS diet until weaning (3 weeks of age) and then randomly assigned to FS or FD diets until 14 weeks of age. (C) Overview of spermatogenesis. Undifferentiated spermatogonia (Undiff) maintain the stem cell population and, upon differentiation cues, give rise to differentiating spermatogonia (Diff). Following meiotic entry, completion of homologous synapsis marks pachytene spermatocytes (PSs), and subsequent reductional divisions generate haploid round spermatids (RSs), which ultimately differentiate into spermatozoa. (D) Experimental design. Testes from adult FS or FD mice were collected for histological analysis or dissociated for fluorescence-activated cell sorting (FACS) of germ cell populations, followed by bulk RNA-seq, ATAC-seq, CUT&Tag, or single-cell RNA-seq. (E) Representative testicular sections from FS (top) and FD (bottom) mice stained with DAPI (left) or immunostained for γH2AX (a marker of PS XY bodies) and H1T (a marker from mid-pachytene onward) (right). Bars, 50 μm. (F) Box-and-whisker plots showing PS density per seminiferous tubule, quantified by γH2AX-positive XY bodies, in FS and FD testes (four biological replicates per condition). Statistical significance was determined using a two-sided Wilcoxon rank-sum (Mann–Whitney U) test. (G) Box-and-whisker plots showing RS density per seminiferous tubule, quantified by H1T-positive cells, in FS and FD testes (three biological replicates per condition). Statistical significance was determined using a two-sided Wilcoxon rank-sum (Mann–Whitney U) test. (H) Transcriptome comparison of FS and FD PSs (three biological replicates per condition). Differentially expressed genes (DEGs; fold change ≥1.5, P < 0.05) are highlighted (red, upregulated in FD; blue, downregulated in FD), with DEG counts indicated. (I) Chromosomal distribution and proportion of CpG island (CGI) genes among PS DEGs. (J) Heat maps showing Z-score–normalized expression of DEGs upregulated (left) or downregulated (right) in FD PSs. THY1^+^, undifferentiated spermatogonia; KIT^+^, differentiating spermatogonia; PS, pachytene spermatocytes; RS, round spermatids.

To assess the impact of folate deficiency on spermatogenesis, we quantified representative germ cell populations in testicular sections from FS and FD mice. Despite dietary folate deficiency, ZBTB16 (PLZF)^+^ undifferentiated spermatogonia—which include the spermatogonial stem cell population— were maintained at similar cell densities in FS and FD testes (Fig. EV1E). Upon commitment to differentiation, KIT^+^ differentiating spermatogonia were also maintained at similar cell densities (Fig. EV1F). These results suggest preservation of mitotically dividing spermatogonia under folate deficiency.

Once germ cells enter meiosis and complete chromosomal synapsis, pachytene spermatocytes (PSs) exhibit unsynapsed sex chromosomes marked by the phosphorylated histone variant H2AX (γH2AX), reflecting activation of the DNA damage response pathway that directs meiotic sex chromosome inactivation (MSCI) and XY body formation (Abe *et al*, 2022; Alavattam *et al*, 2021). Compared with FS testes, FD testes displayed a reduced cell density of PSs exhibiting γH2AX^+^ XY bodies (Fig. 1E, F). In addition to reduced cell density, the relative γH2AX intensity within XY bodies was decreased in FD testes (Fig. EV1G), indicating attenuated MSCI and compromised maintenance of PSs. Accordingly, the overall densities of mid PSs and more advanced germ cells, identified by the testis-specific histone variant H1T, were reduced in FD testes (Fig. 1E, G), whereas SOX9^+^ Sertoli cell densities were comparable between FS and FD mice (Fig. EV1H). Together, these data suggest that spermatogenesis is particularly sensitive to folate deficiency at the pachytene stage.

### Folate deficiency alters transcriptional programs in pachytene spermatocytes

To determine whether folate deficiency perturbs transcription during meiosis, we performed bulk RNA sequencing (RNA-seq) on fluorescence-activated cell sorting (FACS)–purified PSs from FS and FD testes (Fig. 1D). Consistent with prior studies (Alavattam *et al*., 2024; Yeh *et al*, 2021; Yeh *et al*, 2026), FACS isolation yielded highly purified PS populations (Fig. EV2A). RNA-seq biological replicates showed high concordance based on Pearson correlation analysis (Fig. EV2B). Relative to FS controls, FD PSs exhibited 625 upregulated and 545 downregulated differentially expressed genes (DEGs) (Fig. 1H). Of the 625 upregulated DEGs, 68 were located on the sex chromosomes (63 on the X chromosome and 5 on the Y chromosome), including *Rhox11, Cypt1*, and *Pgrmc1*, suggesting perturbation of MSCI (Fig. 1I). Gene ontology (GO) analysis revealed that upregulated DEGs were enriched for spermiogenic pathways, including flagellated sperm motility and spermatid differentiation (Fig. EV1I). These upregulated DEGs are normally highly expressed in wild-type round spermatids (RSs) (Fig. 1J), indicating that folate deficiency induces premature activation of postmeiotic transcriptional programs during meiosis.

In contrast, 99% of downregulated DEGs were encoded on autosomes (Fig. 1I) and included multiple epigenetic regulators and methyltransferases, such as *Dot1l, Mbd3, Mettl14, Prdm2*, and *Kmt2b*. GO analysis of downregulated genes revealed enrichment in chromatin- and RNA-regulatory processes, including mRNA stability and DNA methylation–dependent constitutive heterochromatin formation (Fig. EV1J). More than half of the downregulated DEGs are normally highly expressed during wild-type meiosis (Fig. 1J), indicating perturbation of meiosis-specific transcriptional programs.

In mammals, a large proportion of gene promoters contain CpG islands (CGIs; hereafter referred to as CGI promoters), which are typically maintained in a hypomethylated state (Deaton & Bird, 2011). During spermatogenesis, these hypomethylated CGI promoters are bound by the germline-specific Polycomb protein SCML2, which establishes bivalent chromatin marked by repressive H3K27me3 and active H3K4me3 on thousands of genes in PSs (Maezawa *et al*, 2018a). In contrast, promoters with low CG content are preferentially activated later in haploid RSs (Hammoud *et al*, 2014). Consistent with this regulatory hierarchy, ∼90% of genes downregulated in folate–deficient PSs were CGI-associated (Fig. 1I), indicating that actively transcribed CGI genes during meiosis are particularly sensitive to folate deficiency. Conversely, ∼17% of upregulated genes contained CGIs, consistent with precocious activation of RS-specific transcriptional programs (Fig. 1I). Together, these data demonstrate that folate deficiency impairs CGI-associated meiotic gene regulation and promotes premature postmeiotic gene expression during meiosis.

### Folate deficiency results in loss of active epigenetic modifications in pachytene spermatocytes

Because folate deficiency caused transcriptional dysregulation during meiosis, we next examined whether it alters underlying epigenetic states in meiotic cells. Accordingly, we assessed chromatin accessibility—reflecting regulatory chromatin regions—using assay for transposase-accessible chromatin using sequencing (ATAC-seq). Across five biological replicates of PSs, although some variability among replicates was observed, samples from the FS and FD groups consistently segregated (Fig. EV2C). This pattern indicates a reproducible effect of folate deficiency despite variability among biological replicates, presumably reflecting pleiotropic effects of metabolic perturbation.

Combined analysis of ATAC-seq data across biological replicates identified 1,832 differentially accessible regions (DARs) in FD PSs, including 768 regions with increased accessibility and 1,064 regions with decreased accessibility (Fig. 2A). The majority of DARs are located in intergenic regions or within gene bodies, whereas only a small fraction overlapped transcription start sites (TSSs), suggesting that folate deficiency predominantly affects putative regulatory regions outside of promoters in PSs (Fig. 2B). Notably, gained DARs in FD PSs were enriched on the sex chromosomes, consistent with attenuation of MSCI, whereas most lost DARs were located on autosomes (Fig. 2B).

**Figure 2.**
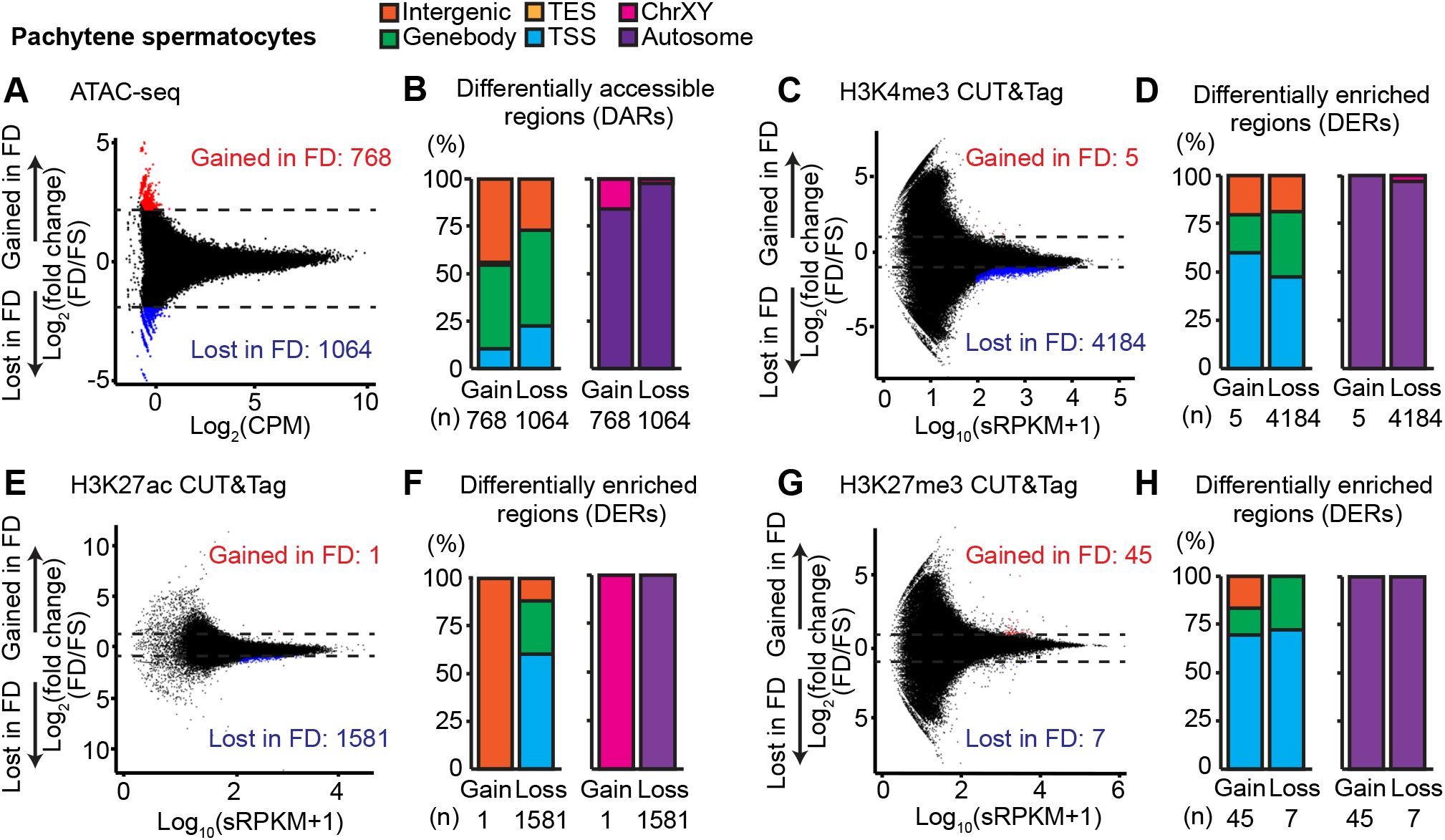
Folate deficiency reshapes chromatin accessibility and histone modification landscapes in pachytene spermatocytes. (A) Comparison of chromatin accessibility in FS and FD PSs (five biological replicates per condition). Differentially accessible regions (DARs; fold change ≥ 2, P < 0.05) are highlighted (red, gained in FD; blue, lost in FD), with counts indicated. (B) Genomic annotation and chromosomal distribution of PS DARs. (C) Comparison of H3K4me3 enrichment in FS and FD PSs (three biological replicates per condition). Differentially enriched regions (DERs; fold change ≥ 1, adjusted P < 0.05) are highlighted (red, gained; blue, lost). (D) Genomic annotation and chromosomal distribution of PS H3K4me3 DERs. (E) Comparison of H3K27ac enrichment in FS and FD PSs (three biological replicates per condition). DERs (fold change ≥ 1, adjusted P < 0.05) are highlighted. (F) Genomic annotation and chromosomal distribution of PS H3K27ac DERs. (G) Comparison of H3K27me3 enrichment in FS and FD PSs (three biological replicates per condition). DERs (fold change ≥ 1, adjusted P < 0.05) are highlighted. (H) Genomic annotation and chromosomal distribution of PS H3K27me3 DERs.

To further examine epigenetic alterations associated with folate deficiency, we analyzed histone modifications using cleavage under targets and tagmentation (CUT&Tag) for the active promoter mark histone H3 lysine 4 trimethylation (H3K4me3), the active promoter/enhancer mark histone H3 lysine 27 acetylation (H3K27ac), and the repressive mark histone H3 lysine 27 trimethylation (H3K27me3). We performed CUT&Tag with spike-in controls normalized for cell numbers to quantitatively evaluate the change in response to folate deficiency. Because H3K4me3 and H3K27me3 are methylation-based modifications requiring SAM, whereas H3K27ac is acetylation-based, this approach allowed us to assess whether methyl-donor limitation selectively disturbs methylation-based marks or more broadly alters histone modifications. For all three assays, samples from three biological replicates in both FS and FD groups consistently segregated by diet (Fig. EV2D). Consistent with the widespread loss of chromatin accessibility, enrichment of the active histone modifications H3K4me3 and H3K27ac was markedly reduced at many loci in FD PSs (Fig. 2C–F). Genomic regions surrounding the *Ddx4* and *Prdm2* loci are shown as representative examples (Fig. EV3A, B). Among differentially enriched regions (DERs) for H3K4me3 and H3K27ac, ∼99% showed decreased enrichment in FD PSs (Fig. 2C, E). In contrast, a modest gain of H3K27me3 DERs was observed in FD PSs (Fig. 2G). Thus, while active chromatin marks were broadly decreased, regions of H3K27me3 enrichment were comparatively less affected, indicating that folate deficiency does not uniformly alter methylation-based histone modifications.

Unlike DARs, which were predominantly located in distal regions, a substantial fraction of DERs for all three histone modifications overlapped TSSs (Fig. 2D, F, and H). Notably, most DERs were located on autosomes, indicating a preferential impact of folate deficiency on autosomal histone modifications compared with sex chromosomes. Together, these data demonstrate that folate deficiency distinctly alters chromatin accessibility and histone modification landscapes during meiosis, with pronounced losses of active histone marks at autosomal promoters.

### Folate deficiency results in loss of active epigenetic modifications on CpG islands

Since we found that actively transcribed CGI genes during meiosis are particularly sensitive to folate deficiency, we next examined how CGI genes are regulated in FD PSs. Active histone modifications, H3K4me3 and H3K27ac, were strongly enriched at CGI gene promoters, which are characterized by hypomethylated DNA, but were markedly reduced at CGI genes in FD PSs (Fig. 3A). In contrast, enrichment of the repressive mark H3K27me3 was increased at CGI genes in FD PSs (Fig. 3A). These results indicate that CGI genes are primary targets of folate deficiency and that loss of folate is associated with reduced H3K4me3 and H3K27ac, thereby counteracting the accumulation of H3K27me3. Consistent with this interpretation, 1,181 and 806 of genes containing lost H3K4me3 and H3K27ac DERs (within the TSS, gene body, or TES) in FD PSs were CGI genes (Fig. 3B), whereas all genes containing ATAC-seq DARs were non-CGI genes. Together, these findings demonstrate that folate deficiency preferentially disrupts histone modifications at CGI genes, while alterations in chromatin accessibility predominantly occur at non-CGI genes.

**Figure 3.**
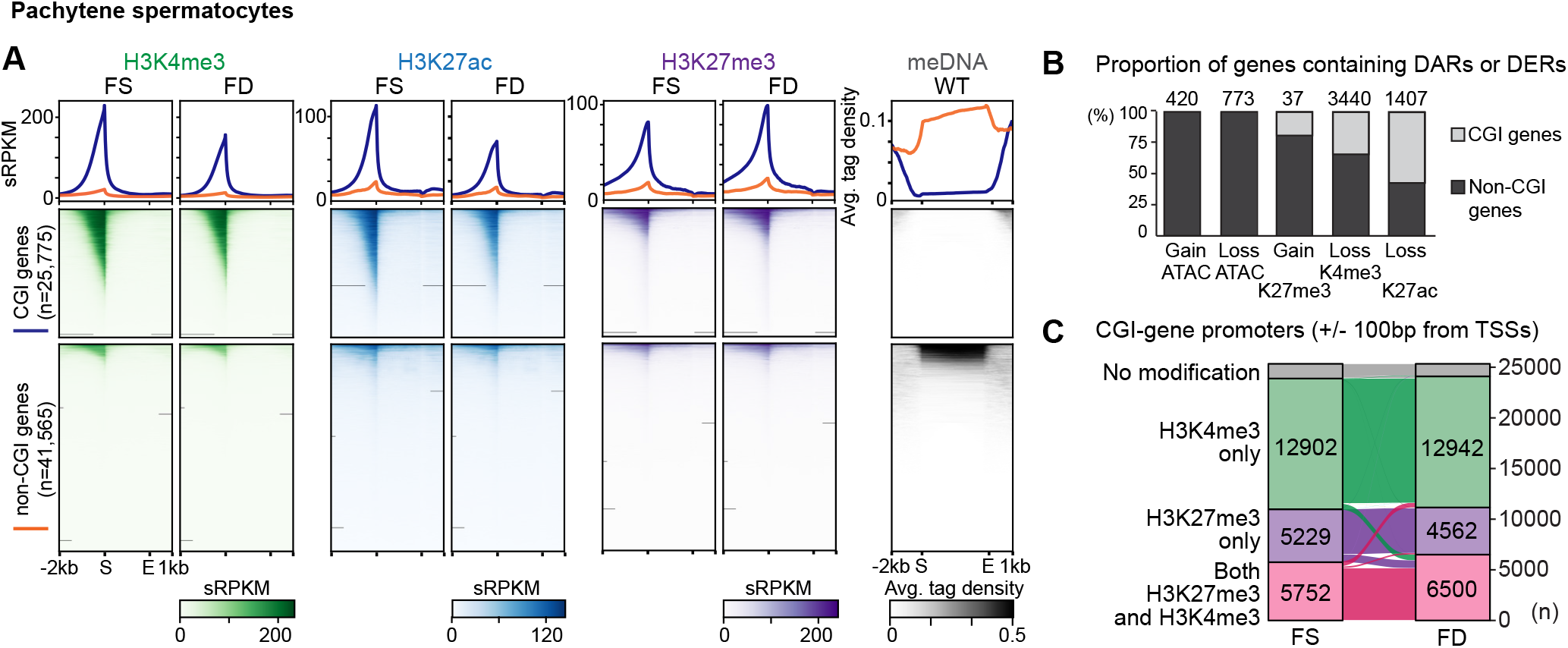
Folate deficiency shifts active and bivalent chromatin states at CpG island promoters in pachytene spermatocytes. (A) CUT&Tag enrichment of H3K4me3, H3K27ac, and H3K27me3 at CGI genes and non-CGI-associated genes in FS and FD PSs. MeDIP-seq enrichment at CGI genes in wild-type PSs was reanalyzed from ref. (Maezawa *et al*., 2025) and is shown for reference. (B) Proportion of PS DARs and DERs (genic regions only) overlapping CGI genes. Counts are indicated. (C) Proportion of CGI promoters (±100 bp from transcription start sites) classified as H3K4me3-only, H3K27me3-only, double-positive, or no modification. Promoter counts are indicated.

At CGI gene promoters, H3K4me3 and H3K27me3 often coexist to form bivalent promoters, a hallmark of PSs that is thought to prime chromatin states for spermiogenesis and subsequent embryogenesis (Lesch *et al*, 2016; Maezawa *et al*., 2018a; Sin *et al*, 2015b). CGI gene promoters (n = 25,775) were classified into four states: H3K4me3 only, H3K27me3 only, both H3K4me3 and H3K27me3 positive, or lacking either modification. When comparing FS and FD conditions, the overall distribution of these states was largely similar; however, the number of CGI gene promoters marked with both H3K4me3 and H3K27me3 increased from 5,752 to 6,500 in FD PSs, arising from promoters previously marked by either H3K4me3 or H3K27me3 alone (Fig. 3C). In contrast, CGI promoters lacking both modifications remained relatively stable between conditions (Fig. 3C). These data further demonstrate that histone modifications at CGI gene promoters are altered in response to folate deficiency.

### CGI genes are subject to epigenomic change in round spermatids in response to folate deficiency

To examine the effects of folate deficiency during postmeiotic haploid stages, we performed bulk RNA-seq on highly purified FACS-isolated RSs from FS and FD testes (Fig. 4A, Fig. EV2A). RS RNA-seq showed strong concordance among biological replicates (Fig. EV2*B*). Relative to FS, FD RS transcriptomes showed minimal changes, with only 21 DEGs—17 upregulated and 4 downregulated—identified (Fig. 4A), indicating that RS transcriptomes are less sensitive to folate deficiency than those of PSs. Notably, all RS DEGs were autosomal, and the majority of upregulated DEGs were classified as CGI genes (Fig. 4B). Despite these modest transcriptional changes, FD RSs exhibited widespread alterations in chromatin accessibility, predominantly on autosomes (Fig. 4C, D). In contrast to PSs, where ATAC DARs were enriched at non-CGI genes, ATAC DARs in RSs were preferentially enriched at CGI genes (Fig. 4E), suggesting that CGI genes are particularly susceptible to chromatin accessibility alteration in RSs in response to folate deficiency.

**Figure 4.**
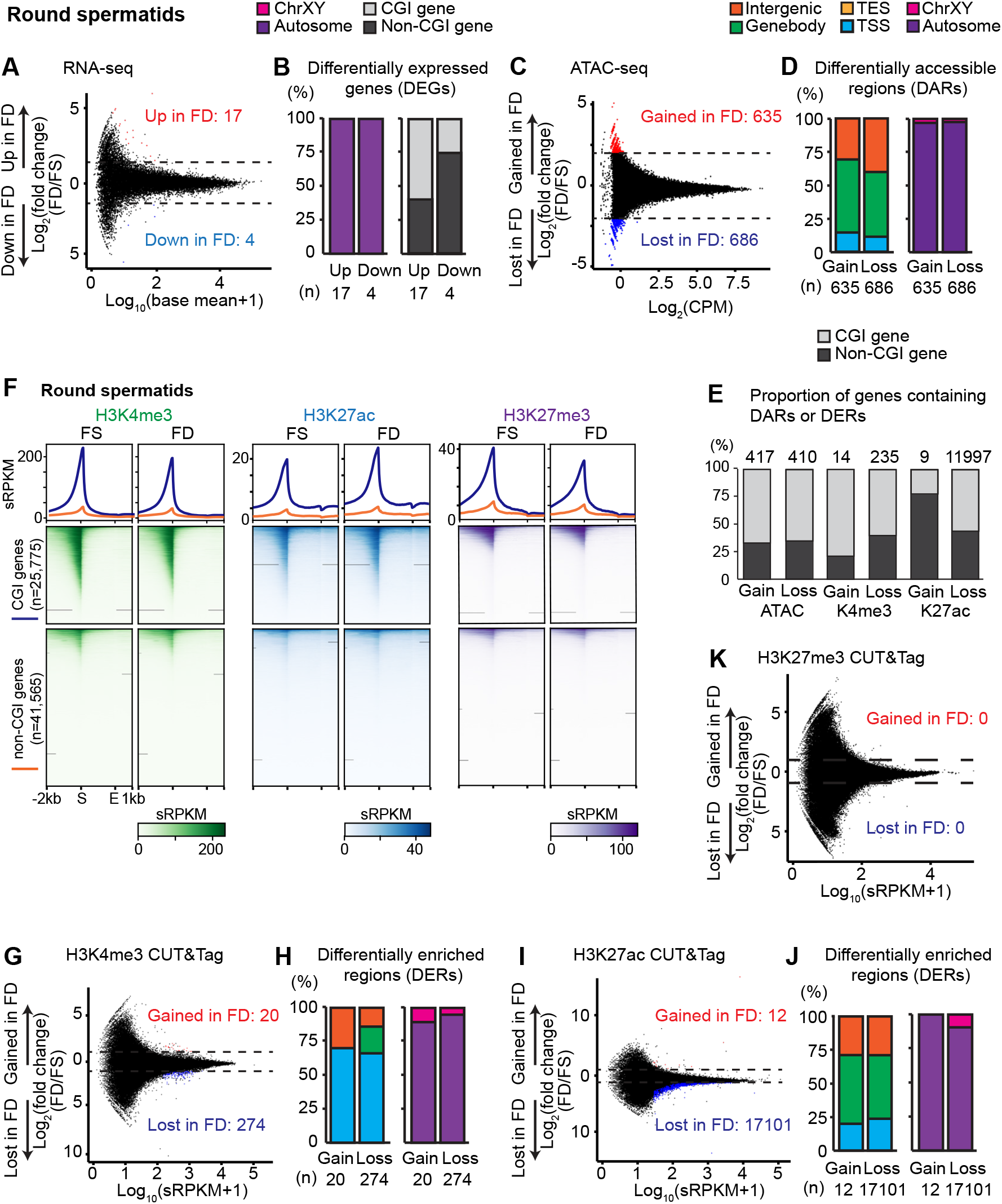
Folate deficiency alters transcriptomic and chromatin landscapes in round spermatids. (A) Transcriptome comparison of FS and FD RSs (three biological replicates per condition). DEGs (fold change ≥ 1.5, P < 0.05) are highlighted. (B) Chromosomal distribution and proportion of CGI genes among RS DEGs. (C) Comparison of chromatin accessibility in FS and FD RSs (five biological replicates per condition). DARs (fold change ≥ 2, P < 0.05) are highlighted. (D) Genomic annotation and chromosomal distribution of RS DARs. (E) Proportion of RS DARs and DERs (genic regions only) overlapping CGI genes. (F) CUT&Tag enrichment of H3K4me3, H3K27ac, and H3K27me3 at CGI genes and non-CGI genes in FS and FD RSs. (G) Comparison of H3K4me3 enrichment in FS and FD RSs (three biological replicates per condition). DERs (fold change ≥ 1, adjusted P < 0.05) are highlighted. (H) Genomic annotation and chromosomal distribution of RS H3K4me3 DERs. (I) Comparison of H3K27ac enrichment in FS and FD RSs (three biological replicates per condition). DERs (fold change ≥ 1, adjusted P < 0.05) are highlighted. (J) Genomic annotation and chromosomal distribution of RS H3K27ac DERs. (K) Comparison of H3K27me3 enrichment in FS and FD RSs. No significant DERs were detected.

Consistent with these findings, FD RSs displayed reduced H3K4me3 intensity at CGI promoters (Fig. 4F) and an overall loss of H3K4me3-enriched regions; 93% of H3K4me3 DERs (274/294) were lost in FD RSs, many of which were located at TSSs (Fig. 4G, H). Notably, 99% of H3K27ac DERs (17,101/17,113) were lost in FD RSs, and the majority of these regions are located in intergenic or gene body regions. In contrast, H3K27ac intensity increased at CGI promoters, suggesting a redistribution of H3K27ac from distal regulatory regions to promoters (Fig. 4F, I, and J). Conversely, H3K27me3 intensity, which counteracts H3K27ac, decreased at CGI promoters without substantial changes in the overall number of H3K27me3 DERs (Fig. 4F, K). Together, these results demonstrate that, following the PS stage, CGI genes continue to exhibit epigenomic alterations in RSs under folate deficiency and suggest that chromatin changes established at CGI genes during PSs, but not those at non-CGI regions, persist into the RS stage.

### Single-cell transcriptome analysis reveals folate–induced adverse effects on spermatogenesis

Having established epigenomic alterations at the bulk-cell level in response to folate deficiency, we next sought to determine how folate deficiency affects transcription at the single-cell level throughout spermatogenesis. To this end, single-cell RNA sequencing (scRNA-seq) was performed on FACS-sorted live testicular cells from two biological replicates of FS and FD mice. The overall cellular composition of the testes was comparable between FS and FD mice (Fig. 5A–C; *Fig. EV*4A). With more than 10,000 reads per cell and a 98.7% mapping rate, the scRNA-seq dataset showed high sequencing quality and detected a total of 17,879 genes (Fig. EV4A, B).

**Figure 5.**
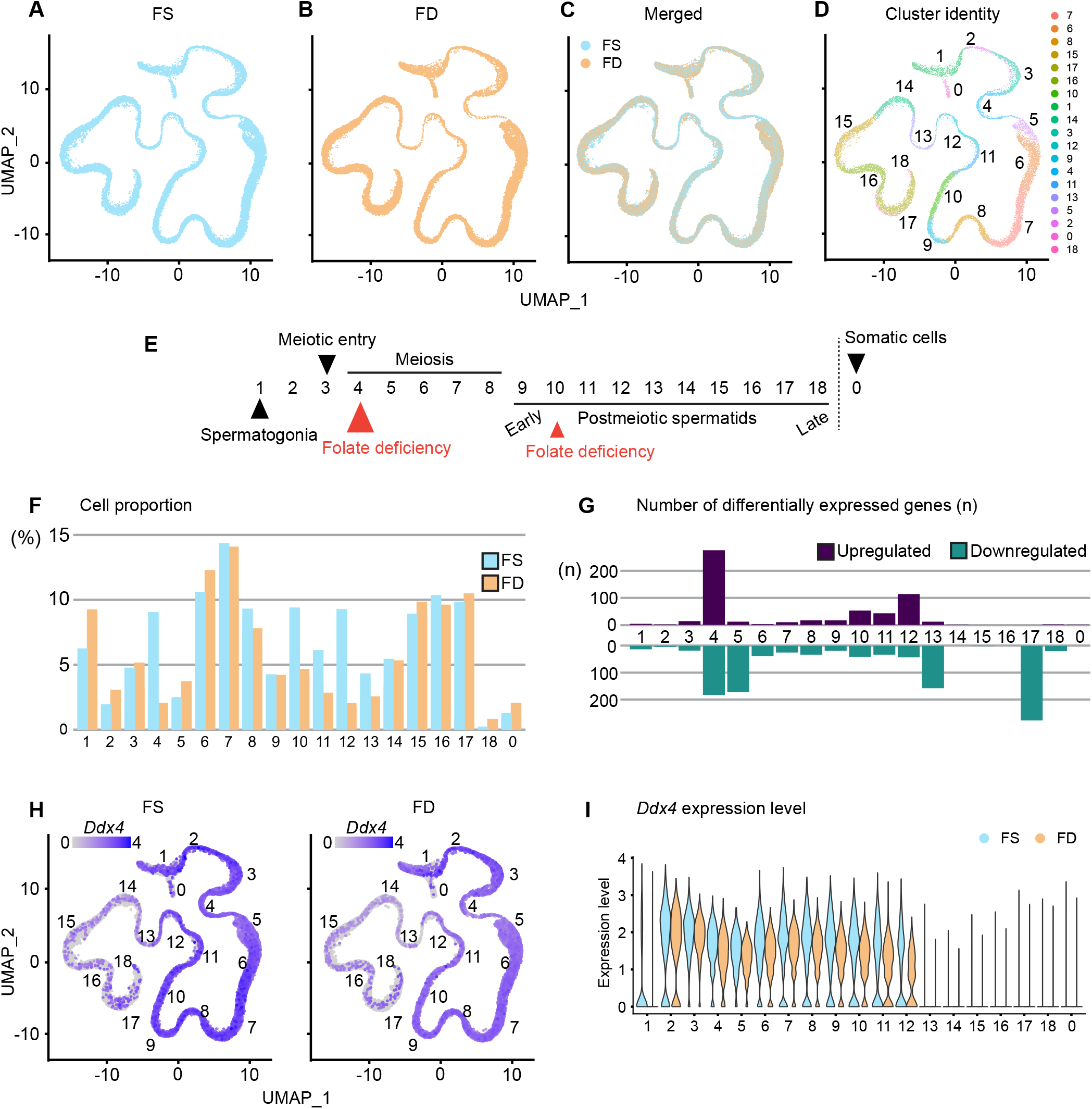
Single-cell transcriptomic profiling reveals altered germ cell composition and cluster-specific transcriptional changes under folate deficiency. (A–C) UMAP projections of germ cell scRNA-seq datasets from FS testes (A), FD testes (B), and integrated FS and FD datasets (C). (D) UMAP showing cluster identity of integrated germ cell datasets. (E) Summary schematic of FD-associated phenotypes across spermatogenic subtypes. Subtype clusters are ordered by inferred developmental progression. Red arrows mark developmental stages where germ cell proportions are substantially reduced in FD mice. (F) Proportion of FS and FD germ cells within each cluster ordered by inferred developmental progression. (G) Number of DEGs per cluster (fold change ≥ 1, adjusted P < 0.05) (purple, upregulated in FD; teal, downregulated in FD), (H) UMAP feature plots showing expression of *Ddx4* in FS and FD datasets. (I) Violin plots showing *Ddx4* expression in FS and FD clusters ordered by inferred developmental progression.

After filtering out low-quality cells and clusters with a high proportion of doublets, uniform manifold approximation and projection (UMAP) analysis identified 19 major cell clusters, and developmental trajectories were inferred based on the expression of key marker genes (Fig. 5D, E; Fig. EV4C–G). Cluster 1 corresponded to spermatogonia, as indicated by high expression of *Kit* (Fig. EV4C). Cluster 4 represented meiotic spermatocytes, consistent with elevated expression of *Pou5f2* (Fig. EV4D). High expression of *Prm1* marked downstream clusters as spermatids (Fig. EV4E). Somatic cells clustered near mitotically dividing spermatogonia and were represented by Cluster 0, as indicated by high expression of *Vim* (Fig. EV4F).

Notably, a germ cell population corresponding to early meiotic spermatocytes (Cluster 4) was markedly reduced in FD testes compared to FS controls (Fig. 5F). In FD testes, reductions in germ cell proportions were also observed in postmeiotic spermatid stages corresponding to Clusters 10–13, representing early to mid postmeiotic spermatids (Fig. 5E, F). Meiotic spermatocytes represented by Clusters 4 and 5 exhibited a higher number of DEGs compared to spermatids in later clusters, whereas fewer DEGs were identified in spermatid clusters (Clusters 10–13) than in spermatocyte clusters (Fig. 5G). These findings demonstrate a major negative impact of folate deficiency on meiotic spermatocytes, consistent with the histological abnormalities observed in FD testes (Fig. 1E-G), with a more modest impact also evident in postmeiotic spermatids.

Interestingly, expression of the germ cell marker *Ddx4*, encoding an essential RNA helicase, was markedly reduced in FD testes (Fig. 5H, I), consistent with reduced enrichment of active histone marks at the *Ddx4* locus (Fig. EV3A). *Ddx4* belongs to a class of germline reprogramming–responsive genes that undergo promoter DNA demethylation in primordial germ cells and play key roles in gametogenesis and meiosis (Hill *et al*., 2018). Reduced expression of several additional germline reprogramming–responsive genes, including *Adad1, Sycp1, Sycp2*, and *Tdrd1*, was observed in FD testes (Fig. EV5A–D). Furthermore, Gene Ontology analysis suggested perturbation of pathways involved in spermatogenesis (Fig. EV4H–K), raising the possibility that disruption of germline regulatory programs contributes to the reduced germ cell density observed in FD testes (Fig. 1E-G).

### H3K4me3 differentially enriched regions in sperm are perturbed earlier in pachytene spermatocytes

Lismer et al. demonstrated that paternal folate deficiency alters H3K4me3 enrichment at developmental loci in mature sperm (Lismer *et al*., 2021). Because these altered H3K4me3 regions are associated with embryonic abnormalities (Lismer *et al*., 2021), we sought to determine whether folate deficiency–induced epigenomic perturbations observed in sperm can be traced to earlier stages of spermatogenesis. Specifically, we examined whether regions exhibiting altered H3K4me3 in FD sperm (Lismer *et al*., 2021) also displayed epigenetic changes in FD PSs and RSs. Strikingly, loci that ultimately become over-enriched for H3K4me3 in FD sperm first exhibited marked depletion of active chromatin marks, including H3K4me3 and H3K27ac, in FD PSs relative to FS controls (Fig. 6A). In FD RSs, H3K4me3 remained reduced at these loci, whereas H3K27ac levels increased (Fig. 6C). Mirroring the H3K27ac profile, H3K27me3 enrichment at the same loci was elevated in FD PSs (Fig. 6A) but reduced in FD RSs compared with FS controls (Fig. 6C). This developmental progression suggests dynamic remodeling of histone marks throughout spermatogenesis in response to folate deficiency.

**Figure 6.**
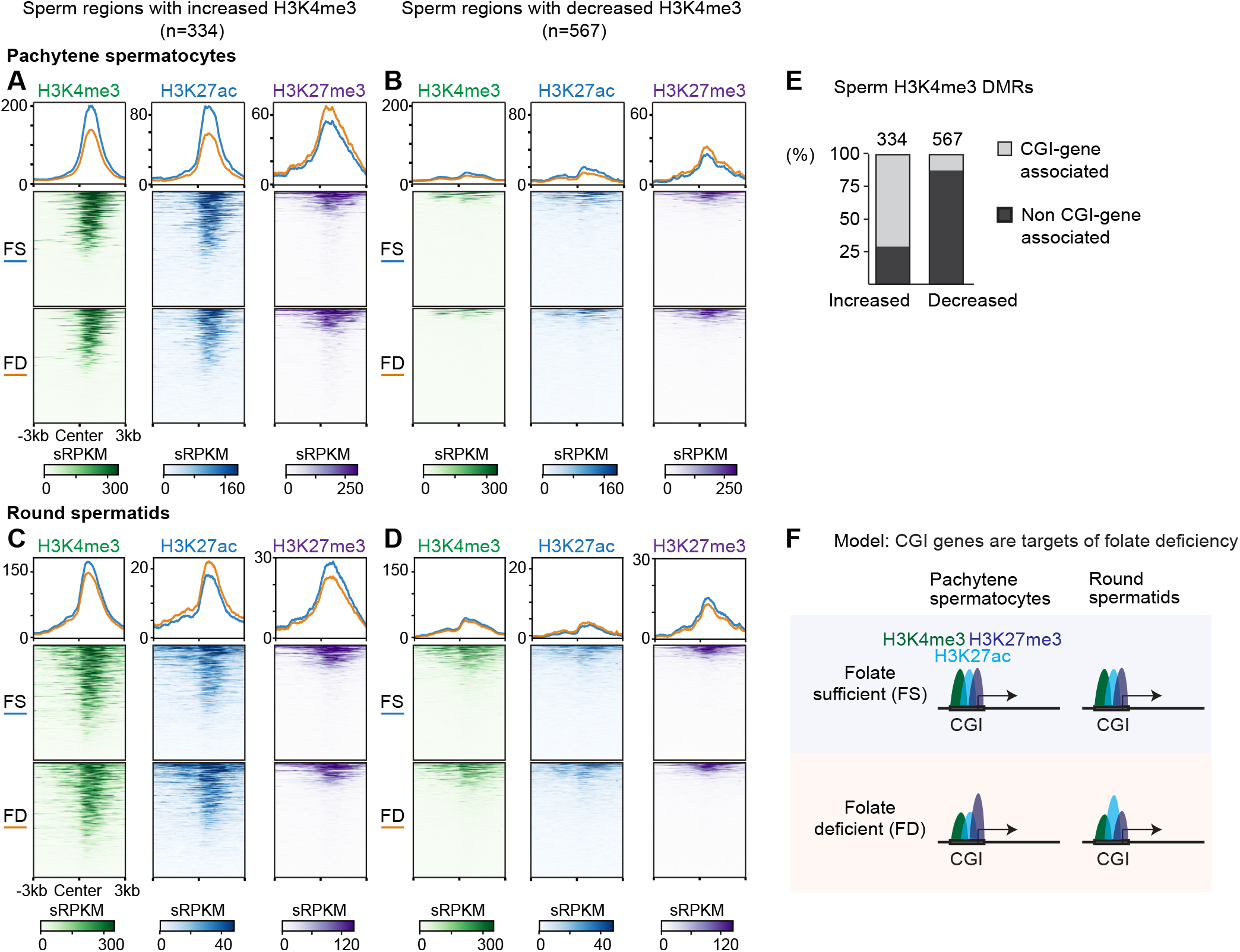
Chromatin regions altered in sperm originate from epigenomic perturbations during meiotic prophase I. (A) CUT&Tag enrichment in FS and FD PSs at regions exhibiting increased H3K4me3 in sperm. (B) CUT&Tag enrichment in FS and FD PSs at regions exhibiting decreased H3K4me3 in sperm. (C) CUT&Tag enrichment in FS and FD RSs at regions exhibiting increased H3K4me3 in sperm. (D) CUT&Tag enrichment in FS and FD RSs at regions exhibiting decreased H3K4me3 in sperm. (E) Proportion of sperm H3K4me3 gain and loss regions overlapping CGI genes. (F) Proposed model of stage-specific chromatin remodeling in response to folate deficiency.

Consistent with Lismer et al. (Lismer *et al*., 2021), 71% of sperm regions showing increased H3K4me3 overlapped CGI genes (Fig. 6E), suggesting that CGI genes represent a chromatin context particularly sensitive to folate-dependent remodeling. In contrast, regions exhibiting decreased H3K4me3 in FD sperm showed minimal epigenomic disruption in PSs or RSs (Fig. 6B, D), and only 11% of these regions overlapped CGI genes (Fig. 6E). Together, these findings indicate that gains and losses of H3K4me3 in FD sperm arise from distinct genomic contexts, with CGI genes exhibiting heightened developmental responsiveness to folate deficiency.

## Discussion

Epigenetic programming during spermatogenesis is essential for proper sperm formation and for supporting normal offspring development. Although paternal nutrition has emerged as an important determinant of offspring health, how metabolic perturbations influence epigenetic regulation across spermatogenic stages has remained unclear. Previous studies showed that folate deficiency alters H3K4me3 patterns in mature sperm, suggesting sensitivity of the sperm epigenome to methyl-donor availability. Here, we demonstrate that paternal folate deficiency alters spermatogenesis beginning at meiotic prophase I, affecting both transcriptional programs and chromatin organization. These findings reveal that epigenetic perturbation originates during meiosis rather than arising solely during late spermiogenesis and highlight the substantial metabolic and epigenetic demands required to sustain gene regulation in PSs. Together, our findings identify meiotic prophase I as a metabolically sensitive window that integrates paternal nutritional status into the germline epigenome. In this framework, meiosis functions as a developmental interface through which metabolic conditions can be translated into stable epigenetic states in the male germline.

In 1984, Robin Holliday proposed that meiosis functions not only to generate haploid gametes and promote genetic recombination but also to reset epigenetic information for the next generation (Berger, 2024; Holliday, 1984). Although this concept has long been neglected, recent studies have revealed that spermatogenesis is accompanied by extensive chromatin reorganization during the transition from mitotically dividing spermatogonia to meiotic spermatocytes, including large-scale transcriptional activation, histone modification remodeling, and chromatin structural reconfiguration (Alavattam *et al*., 2024; Hasegawa *et al*., 2015; Kitamura *et al*., 2025a; Kitamura & Namekawa, 2024, 2025; Kitamura *et al*., 2025b; Lin *et al*., 2025; Maezawa *et al*., 2020b; Maezawa *et al*., 2018b; Maezawa *et al*., 2021; Rabbani *et al*., 2022; Sin *et al*., 2015a). Our findings extend this concept by suggesting that meiotic prophase I operates as a metabolic sensing window in the male germline, during which environmental and nutritional conditions are integrated into the epigenomic landscape undergoing reprogramming. This stage-specific sensitivity provides a mechanistic framework for understanding how paternal nutritional status can shape heritable chromatin states in sperm.

Unlike somatic tissues, spermatogenesis represents a lifelong maintenance system sustained by a preserved spermatogonial stem cell population. Concentrating environmentally responsive epigenetic remodeling during meiosis may therefore represent a biological strategy that allows transient metabolic perturbations to influence germ cell epigenetic states without compromising stem cell integrity. Such stage-restricted responsiveness enables temporal environmental information to be encoded while safeguarding long-term germline continuity despite fluctuating metabolic conditions throughout adult life. In this framework, meiotic prophase I functions as a developmental buffering stage that balances environmental responsiveness with germline stability. This model provides a mechanistic explanation for how adult paternal exposures can influence sperm epigenomic states while preserving sustained fertility.

The epigenomic response to folate deficiency was not uniform across spermatogenesis. Active chromatin marks, including H3K4me3 and H3K27ac, were reduced at CGI promoters in FD PSs, whereas the repressive mark H3K27me3 was increased at these promoters (Fig. 3 and 6F). Because both H3K4me3 and H3K27me3 depend on SAM as a methyl donor, the differential sensitivity of these marks suggests selective prioritization or stabilization of Polycomb-associated repression under conditions of methyl-donor limitation. These observations indicate that active chromatin states are particularly vulnerable to metabolic stress, whereas repressive chromatin programs may be comparatively protected during meiosis.

Our analyses further identify CGI genes as primary epigenomic sites of vulnerability under folate deficiency. CGI promoters are typically hypomethylated and enriched for bivalent chromatin in PSs, a configuration thought to poise developmental regulators for precise activation or repression during spermatogenesis and early embryogenesis (Lesch *et al*., 2016; Maezawa *et al*., 2018a; Sin *et al*., 2015b). Under folate deficiency, we observed widespread depletion of active histone marks at CGI genes accompanied by redistribution toward increased bivalency or relative H3K27me3 dominance. Notably, CGI loci that ultimately acquire strong H3K4me3 enrichment in mature sperm exhibited marked depletion during meiosis, indicating that folate-dependent chromatin remodeling at these promoters is dynamic and developmentally progressive. Our analysis integrates previously published sperm epigenomic datasets with stage-resolved profiling of germ cells to infer the developmental origin of sperm chromatin alterations. Together, these findings support a model in which methyl-donor limitation perturbs the balance between active and repressive histone modifications during meiosis, potentially predisposing specific loci to altered epigenetic states in sperm.

These observations further suggest that folate-dependent chromatin remodeling during meiosis may intersect with additional layers of chromatin regulation. Recent work has shown that histone variant exchange mediated by the chaperone DAXX promotes replacement of testis-specific histone H3 variants with H3.3 during meiotic prophase I (Yeh *et al*., 2026). Because H3.3 preferentially marks retained nucleosomes in mature sperm and bivalent chromatin domains are implicated in paternal epigenetic inheritance (Erkek *et al*, 2013; Sakashita *et al*, 2023), our findings raise the possibility that folate-dependent chromatin regulation intersects with histone variant dynamics. This perspective suggests that metabolic perturbation may influence multiple layers of germline chromatin organization beyond individual histone modifications, contributing to long-lasting epigenomic consequences.

The sensitivity of CGI genes to folate deficiency further raises the possibility that maintenance of DNA hypomethylated states may become destabilized under methyl-donor limitation. Although genome-wide DNA methylation remains largely stable during spermatogenesis, our recent study has revealed locus-specific methylation changes (Maezawa *et al*, 2025). Whether CGI promoters maintain methylation fidelity under conditions of limited methyl-donor availability remains unresolved. Future studies integrating histone modification, histone variant dynamics, chromatin accessibility, and DNA methylation will be necessary to define how metabolic stress reshapes multilayered epigenetic regulation in the germline and how these changes influence paternal epigenetic inheritance.

Together, our findings identify meiotic prophase I as a metabolically sensitive stage that integrates paternal nutritional status into the germline epigenome. Rather than passively reflecting environmental stress, the meiotic program appears positioned to selectively encode metabolic information during a developmental window characterized by extensive chromatin remodeling and transcriptional reorganization. This stage-specific responsiveness provides a conceptual framework explaining how adult environmental exposures can influence offspring phenotypes without compromising long-term germline stability maintained by spermatogonial stem cells. More broadly, these results support a model in which meiosis functions as an adaptive interface between environment and heredity, translating transient metabolic conditions into persistent epigenetic states transmitted through sperm. Defining how metabolic inputs intersect with chromatin regulation during meiosis will be essential for understanding the molecular basis of paternal epigenetic inheritance and how environmental factors shape developmental trajectories across generations.

## Methods

### Animal model and dietary treatment

Male C57BL/6J mice were maintained under standard conditions and randomly assigned to FS or FD diets after weaning. Mice remained on their assigned diets for 11 weeks and were analyzed at 14 weeks of age. All procedures were approved by the Institutional Animal Care and Use Committee at the University of California, Davis. Detailed experimental conditions are provided in the Supplementary Method.

### Histology and germ cell isolation

Testes were processed for histological and immunofluorescence analyses using established procedures. PSs and RSs were isolated from dissociated testes by FACS as previously described. Detailed protocols are provided in the Supplementary Method.

### Next-generation sequencing and computational analysis

RNA-seq, ATAC-seq, CUT&Tag, and scRNA-seq were performed on purified germ cell populations. Libraries were sequenced on Illumina platforms and analyzed using standard bioinformatic pipelines. Differential gene expression and chromatin analyses were performed using established statistical frameworks. Detailed experimental procedures, sequencing parameters, and computational analyses are described in the Supplementary Method.

### Statistical analysis

Statistical analyses were performed using R. Replicate numbers and statistical tests are indicated in the figure legends and described in the Supplementary Method.

## Data availability

The accession codes for the raw published data used in this study are as follows:

- THY1^+^, PS, and RS RNA-seq (Fig. 1J): GSE55060 (Hasegawa *et al*., 2015)
- KIT^+^ RNA-seq (Fig. 1J): GSE89502 (Maezawa *et al*., 2018a)
- MethylCap-seq (Fig. 3A) GSE262669 (Maezawa *et al*., 2025)
- Sperm H3K4me3 (Fig. 6A-D) GSE135678 (Lismer *et al*., 2021)

Sperm H3K4me3 data (GSE135678) (Lismer *et al*., 2021) were reanalyzed, and differentially enriched regions (DERs) were identified using the following criteria: a p-value < 0.1 (quasi-likelihood F-test) and an average log2 CPM > 1.

## Acknowledgments

We thank members of the Namekawa lab for discussions; AI Ikuyo for discussions and help with data analysis; and Mary Ann Handel for providing the H1T antibody. We acknowledge the following funding sources: the NIGMS T32 predoctoral training program in Molecular and Cellular Biology (MCB) at UC Davis (T32GM007377) and a National Academies Ford Foundation Fellowship to J.M.E.; UC Davis Provost’s Undergraduate Fellowship to R.V.K.; R35GM141085 to S.H.N.; and a supplement to R35GM141085 to J.M.E. and S.H.N.

## Author contribution

J.M.E. and S.H.N. designed the study. J.M.E., R.V.K., and S.J. performed the experiments, with contributions from M.H. and Y.M. J.M.E. performed the computational analysis, with contributions from M.H., Y.M., A.S., and S.M. J.M.E., A.S., S.M., R.M.S., and S.H.N. interpreted the results. J.M.E. and S.H.N. wrote the manuscript with critical feedback from all authors. S.H.N. supervised the project.

