## Supplemental Material for "Paternal folate deficiency reveals meiosis as a metabolic sensing window in the male germline"

**Supplemental methods**

**Animals**

Mice were maintained on a 12:12 light:dark cycle in a temperature- and humidity-controlled vivarium (22 ± 2˚C; 40-50% humidity) with *ad libitum* access to food and water in a pathogen-free animal care facility. All animal procedures were approved by the Institutional Animal Care and Use Committee (IACUC protocol no. 21931 and 23545) at the University of California, Davis. All mice were maintained on a C57BL6/J background (Charles River Laboratories).

**Dietary treatments**

Dietary folate concentrations were based on established paternal diet mouse models (1, 2). The folate sufficient (FS) diet contained 2.0 mg/kg folic acid (TD.01369, Inotiv), whereas the folate deficient (FD) contained 0.3 mg/kg folic acid (TD.01546, Inotiv). All breeder mice were maintained on the FS diet. Male littermates were weaned at 3 weeks of age, and litters with at least two males were randomly assigned to either the FS or FD diet. Mice remained on their assigned diets for 11 weeks and were sacrificed at 14 weeks of age.

**Preparation of meiotic chromosome spreads**

Meiotic chromosome spreads were prepared as previously described (3). Testes were excised, detunicated, and seminiferous tubules were detangled in 1x phosphate-buffered saline (PBS). Tubules were sequentially washed in PBS and incubated in hypotonic extraction buffer (HEB) (30 mM Tris base, 17 mM trisodium citrate, 5 mM ethylenediaminetetraacetic acid (EDTA), 50 mM sucrose, 5 mM dithiothreitol (DTT), and 1x cOmplete Protease Inhibitor Cocktail (Sigma, 11836145001) on ice for 1 hour. Tubules were mechanically dissociated in 100 mM sucrose, and cell suspensions were applied to positively charged slides (Probe On Plus, Thermo Fisher Scientific, 22-230-900) precoated with fixation solution (2% paraformaldehyde, 0.1% Triton X-100, and 0.02% sodium monododecyl sulfate, adjusted to pH 9.2 with sodium borate buffer). Slides were incubated in a humid chamber at room temperature for a minimum of 4 hours, air-dried, washed twice in 0.4% Photo-Flo (Kodak, 146-4510) and stored at -80˚C until use.

**Histology and Immunostaining**

Testes were fixed with 4% paraformaldehyde with 0.1% Triton X-100 in PBS at 4˚C overnight, dehydrated in 70% ethanol, and embedded in paraffin. Sections (4 μm) were deparaffinized and subjected to antigen retrieval in citrate buffer (pH 6.1, DAKO, S-1700) using an autoclave (121˚C, 15 psi, 10 minutes). Sections were blocked in Blocking One Histo (Nacalai USA, 06349-64) and then incubated in primary antibodies at 4˚C overnight. Alexa Fluor-conjugated secondary antibodies were applied for 1 hour at room temperature. Slides were counterstained with DAPI and mounted in ProLong Gold. Images were obtained with a Nikon AX Confocal microscope system and were processed using NIS-Elements Advanced Research (Nikon), Photoshop (Adobe), and Illustrator (Adobe) software. Images were acquired using identical microscope settings (laser power, gain, and exposure time) across experimental conditions within each staining experiment.

This study used the following primary antibodies at the following dilutions [format: host-protein (source or company with product/catalog number if applicable), dilution]: goat anti-ZBTB16/PLZF (R&D Systems, AF2944), 1/200; goat anti-cKIT (R&D Systems, AF1356), 1/500; mouse anti-H2AX-pS139 conjugated Alexa 647 (γH2AX) (Millipore, 05-636), 1/500; guinea pig anti-H1T (a gift from Dr. Mary Ann Handel (4)), 1/500; rabbit anti-SOX9 (Millipore, AB5535), 1/500).

**FACS-isolation of male germ cells**

PSs, and RSs were isolated using Fluorescence-activated cell sorting (FACS) as described previously (5). In brief, single cell suspensions were generated from seminiferous tubules by enzymatic dissociation in DMEM containing 2mg/ml Type 1 collagenase (Worthington, CLS1), 0.25 mg/ml DNase I (Sigma, D5025), 1.5 mg/ml hyaluronidase (Sigma, H3506), and 700 U/ml recombinant collagenase (FUJIFILM Wako Pure Chemical Corporation, 036-23141) at 35˚C followed by several PBS washes. Cells were filtered through 70 μm strainers (Falcon, 352350) and washed in FACS buffer (PBS + 2% FBS). PSs and RS were isolated using 2 μl/ml Vybrant DyeCycle Violet. Sorting was performed on a Cell Sorter SH800 (SONY) into collection buffer (50% FBS diluted in PBS). After sorting, cells were washed in PBS, resuspended in CELLBANKER 1 cryopreservation medium (Zenogen Pharma Co, 11910), and stored at -80˚C until use.

**Bulk RNA-seq library generation and sequencing**

FACS-isolated PSs (~50,000) and RSs (~100,000) were collected from three independent biological replicates. ERCC RNA Spike-In Mix 1 (Invitrogen, 4456653) diluted 1:10,000 was added prior to RNA extraction (5μl for PS samples, 10μl for RS samples). Total RNA was isolated using the RNeasy Plus Mini Kit (QIAGEN, 74134) according to the manufacturer's instructions. Libraries were prepared using the NEBNext® Single Cell/Low Input RNA Library Prep Kit for Illumina® (NEB, E6420S) sequenced on the Illumina NovaSeq X Plus platform (150 bp paired-end).

**Bulk RNA-seq read processing, quantification, and differential expression analysis**
Paired-end RNA-seq reads were adapter- and quality-trimmed using Trim Galore (https://github.com/FelixKrueger/TrimGalore) with parameters --paired --nextera --quality 25 --length 30 and FastQC reporting. Trimmed reads were aligned to mm10 (GRCm38) using HISAT2 (v2.2.1). Alignments were converted to BAM format using SAMtools (v1.16.1), and uniquely mapped reads were retained (MAPQ ≥30; flags -q 30 -F 4). Filtered BAM files were sorted and indexed.

Gene-level counts were generated using featureCounts (Subread v2.0.2) with the GENCODE vM25 annotation in paired-end mode (-p -B -C). Counts were generated separately for each biological replicate.

Differential expression analysis was performed using DESeq2 (v1.38.3). For each cell type (PS and RS), count matrices were constructed by merging replicate count files by gene identifier. The design formula was: ~ batch + condition where batch corresponded to biological replicate. Differential expression was assessed using the Wald test. Differentially expressed genes (DEGs) were defined using nominal P value < 0.05 and |log2FoldChange| > 1.5.

For quality control, variance-stabilizing transformation (VST; blind = TRUE) was applied. Pearson correlation matrices were calculated from VST-transformed counts and visualized using pheatmap. MA plots were generated using ggplot2 (v3.5.1).

**ATAC-seq library preparation and sequencing**
ATAC-seq libraries were prepared as previously described from FACS-isolated PSs (~50,000) and RSs (~100,000) collected from five independent biological replicates. Nuclei were isolated and subjected to Tn5-transposase (made in-house) at 37˚C for 30 min. After tagmentation, the transposed DNA was purified using the MinElute kit (Qiagen) following the manufacturer's instructions. Libraries were amplified using PCR, with cycle number determined by qPCR to reach 25% saturation. Libraries were purified using SPRIselect beads (Beckman Coulter) and sequenced on an Illumina NovaSeq X Plus (150-bp paired end).

**ATAC-sequencing read processing, peak calling, and differentially accessibility analysis**
Raw paired-end ATAC-seq reads were adapter- and quality-trimmed using Trimgalore (v 0.6.10) with FastQC reporting. Trimmed reads were aligned to the mouse reference genome (mm10/GRCm38) in paired-end mode with parameters –end-to-end –no-mixed –no-discordant -I 10 -X 200. Alignments were converted to BAM format, sorted, and indexed using SAMtools (v 1.16.1).

To account for Tn5 insertion offsets, alignments were shifted using deepTools alignmentSieve (--ATACshift; v3.5.2). Uniquely mapped, properly paired reads were retained (SAMtools flags -f 2 -F 2048, MAPQ ≥30), and mitochondrial reads (chrM) were removed. ENCODE mm10 blacklisted regions (v2) were excluded using BEDTools (v2.31.0). Library complexity was assessed using csaw (readsDupFreq and estimateLibComplexity; v1.32.0). To normalize libraries to a common molecular complexity following the approach of Reske *et al*. (6), BAM files were downsampled using SAMtools to match the lowest number of unique fragments across the experiments per cell type. PCR duplicates were removed using Picard MarkDuplicates (v 3.0.0) (https://broadinstitute.github.io/picard/) with REMOVE_DUPLICATES=true, and final BAM files were sorted and indexed.

Peaks were called on each biological replicate using MACS3 (v3.0.0b3) with parameters –nomodel –nolambda -q 0.05 -g mm -B –SPMR –keep-dup all. Peak sets were filtered to remove blacklist region overlaps and non-standard contigs. Condition-level peak sets were obtained by pooling replicate BAMs within each condition and calling peaks with the same MACS3 parameters. A consensus peak file was defined as the union of the pooled peak sets from FS and FD samples.

Differential accessibility was assessed in R using csaw and edgeR (v3.40.2) using the consensus peak file as the working feature set ("peaks-only" analysis). Read counts over the consensus peaks were computed with csaw regionCounts using paired-end mode, restricting to standard chromosomes (chr1-19, X, Y), and discarding blacklisted regions. Low abundance peaks were filtered by average abundance (edgeR avgLogCPM threshold > -3). Library size normalization factors were computed using TMM normalization from binned genomic windows (csaw windowCounts with 10-kb bins) and applied to peak counts (csaw normFactors), following Reske *et al.* Differential accessibility was tested using edgeR quasi-likelihood negative binomial generalized linear models (estimateDisp; glmQLFit with robust=TRUE; glmQLFTest) comparing FD against FS samples.

To report region-level differential accessibility regions (DARs), nearby significant windows were merged (csaw mergeWindows; tolerance 500bp; maximum merged width 5kb), and each merged region was assigned statistical significance based on its most significant constituent window (csaw getBestTest). DARs were defined using P-value threshold of <0.05 and an absolute log2 fold-change threshold of ≥ 2. Normalized coverage tracks were generated using deepTools bamCoverage with BPM normalization.

**CUT&Tag library preparation and sequencing**
FACS-isolated PSs (~50,000) and RSs (~100,000) were collected from three independent biological replicates. CUT&Tag libraries of PSs and RSs were prepared as previously described (7, 8) with some modifications (a step-by-step protocol https://www.protocols.io/view/bench-top-cut-amp-tag-kqdg34qdpl25/v3) using CUTANA™ pAG-Tn5 (Epicypher, 15-1017). To perform quantitative spike-in CUT&Tag, Drosophila S2 cells were added to mouse germ cells at a fixed ratio (1000 S2 cells to 5000 mouse germ cells) at the beginning of each reaction. The antibodies used were rabbit anti-H3K4me3 (1/50; Invitrogen; 703849, LOT 2920880), rabbit anti-H3K27ac (1/50; Cell Signaling Technology; 8173S), and rabbit anti-H3K27me3 (1/50; Cell Signaling Technology; 9733S). CUT&Tag libraries were sequenced on Illumina NovaSeq X Plus (150-bp paired end).

**CUT&Tag data processing, spike-in normalization, and differential enrichment analysis**
CUT&Tag paired-end reads were processed following the Henikoff Lab pipeline with modifications. Reads were adapter- and quality-trimmed using Trim Galore (v0.6.10) (https://github.com/FelixKrueger/TrimGalore) and aligned to mm10 (GRCm38) using Bowtie2 (v2.4.5) with parameters --end-to-end --very-sensitive --no-mixed --no-discordant --phred33 -I 10 -X 700. Spike-in Drosophila melanogaster DNA (S2-derived) was aligned separately to dm6 using Bowtie2 with additional parameters --no-overlap --no-dovetail to prevent cross-mapping.

Mouse-aligned BAM files were sorted and indexed using SAMtools (v1.16.1). ENCODE mm10 blacklist regions (v2) were removed using BEDTools (v2.31.0), and properly paired high-confidence reads were retained (SAM flags -f 2 -F 2048, MAPQ ≥30). Library complexity was estimated using ATACseqQC (readsDupFreq, estimateLibComplexity), and within each cell type × histone mark group, BAM files were downsampled to the minimum unique fragment count across replicates. PCR duplicates were removed using Picard MarkDuplicates (v3.0.0) (https://broadinstitute.github.io/picard/).

Spike-in normalization factors were calculated per replicate as: scale factor = 100,000 / (aligned dm6 reads)
Replicate-level spike-in–scaled coverage tracks were generated using deepTools bamCoverage (v3.5.5) with parameters --binSize 1 --extendReads --samFlagInclude 64 --normalizeUsing RPKM --scaleFactor <sample-specific factor>.
For visualization only, biological replicates were merged using SAMtools merge. Group-level scale factors were calculated as: scale factor = 100,000 / (sum of aligned dm6 reads across replicates). Merged spike-in–scaled BigWig tracks were generated for genome browser visualization (IGV v2.14.1).

Replicate concordance was assessed using deepTools multiBigwigSummary (bins mode). Log2-transformed signals were batch-corrected using limma (removeBatchEffect), and Pearson correlation matrices were visualized using pheatmap.

For differential enrichment analysis, replicate-level scaled BigWig files were summarized using multiBigwigSummary (bins mode). The resulting matrices were analyzed using DESeq2 (v1.38.3). Because coverage tracks were spike-in normalized prior to modeling, size factors were fixed to 1. Region values were scaled by a constant and rounded to integers prior to modeling. Differential enrichment was assessed using the design formula ~ batch + condition, comparing FD and FS samples using the Wald test. Differentially enriched regions (DERs) were defined using adjusted P value < 0.05 and |log2FoldChange| > 1.

**Single-cell RNA-seq library preparation**
Single cell suspensions were prepared from testes collected from two independent biological replicates as described above. Dead cells were excluded by 7-AAD Viability Stain (35 µl/ml), and viable cells were sorted prior to fixation. For each biological replicate, at least 1 x $10^6$ viable testicular cells were collected. Cells were fixed for 22 hours at 20 ˚C using the Chromium Next GEM Single Cell Fixed RNA Sample Preparation kit (10X Genomics, 1000414). Approximately 20,000 cells were loaded on the Chromium Controller (10X Genomics Inc.). Libraries were prepared using the Chromium

GEM-X Flex Reagent kits (CG000787 Rev A) and sequenced on the Element Biosciences AVITI system using a paired-end read configuration of 28 bp (Read 1) × 10 bp (i7 index) × 10 bp (i5 index) × 90 bp (Read 2).

**Single-cell RNA-seq processing and analysis**

Alignment and gene counting was performed using the cloud-based 10x Genomics Cell Ranger Multi pipeline (v9.0.0) against the mouse reference genome mm10 (GRCm39) 2024-A.

Downstream analysis was performed in R using Seurat (v5.3.0; SeuratObject v5.1.0). Feature-barcode matrices were loaded using Read10X_h5 and converted to Seurat objects with an initial filter of ≥200 detected genes per cell. The fraction of mitochondrial transcripts was calculated using PercentageFeatureSet (pattern "^mt-"), and low-quality cells were removed by filtering for 200 < nFeature_RNA ≤ 4,000 and mitochondrial content <10%. Each sample was normalized using SCTransform (SCTransform, default parameters), and datasets were integrated using SCT-based integration with default integration features (SelectIntegrationFeatures), anchor identification (FindIntegrationAnchors, normalization.method = "SCT"), and data integration (IntegrateData). Principal component analysis was performed on the integrated assay (RunPCA), and clustering was carried out using a shared nearest-neighbor graph constructed from the first 10 principal components (FindNeighbors, dims = 1:10) followed by Louvain clustering (FindClusters, resolution = 0.4). UMAP embeddings were computed using the first 10 principal components (RunUMAP, dims = 1:10, min.dist = 0.3, spread = 1).

Doublets were identified using scDblFinder (v1.20.2) by converting the integrated Seurat object to a SingleCellExperiment and running scDblFinder. Cells classified as singlets were retained for downstream analyses. Clusters enriched for predicted doublets were identified by inspection of scDblFinder classifications and marker expression patterns and were excluded prior to re-running scaling, PCA, neighbor finding, clustering (resolution = 0.4), and UMAP on the curated dataset.

For cluster-resolved differential expression testing between dietary conditions, the RNA assay was used. RNA layers were joined (JoinLayers) and normalized log-expression values were computed with NormalizeData (RNA assay). Within each cluster, cells were subset and identities were set to dietary group; differential expression between folate-deficient and folate-sufficient samples was tested using FindMarkers (Seurat default test) on the RNA assay using the normalized data layer (layer = "data"), with only.pos = FALSE. Genes were considered differentially expressed using thresholds of adjusted P value <0.05 and |log2 fold change| ≥ 1.

**Identification of CGI genes and non-CGI genes**

CpG islands (CGIs) were defined as published (9, 10), and their coordinates of mouse genome GRCm38/mm10 assembly (n=17,017) were downloaded from UCSC genome browser (https://genome.ucsc.edu/cgi-bin/hgTables?db=mm10&hgta_group=regulation&hgta_track=cpgIslandExt&hgta_table=cpgIslandExt&hgta_doSchema=describe+table+schema). Genes were classified as CGI genes if a first base pair of any transcript TSSs ±100-bp overlapped a CGI. In this way, the total 67,340 genes from Gencode vM25 mouse gene annotations (after removing Chromosome M genes) were divided into CGI genes (n=25,775) and non-CGI genes (n=41,565).

**Quantification and statistical analysis**

All image quantifications were performed in a blinded manner. Images were exported as individual experimental units (e.g. individual seminiferous tubules for histological analysis) and file names were anonymized prior to analysis using the Blind Analysis Tool plugin for ImageJ. Investigators performing quantifications were unaware of dietary group assignments, and encrypted file names were decoded only after completion of analysis for statistical analysis. Signal intensity measurements were performed using NIS Elements software. Fluorescence intensity values were normalized to DAPI signal to control for variation in nuclear content and imaging conditions. For tubule-based quantifications, the number of positive cells per tubule was determined and normalized to tubule surface area, which was measured using manual region-of-interest tracing in NIS-Elements.

**Statistics**

Statistical methods and P values for each plot are listed in the figure legends and/or in the Methods. Statistical significance for pairwise comparisons was determined using two-tailed unpaired t-tests. Next-generation sequencing data were based on at least three independent replicates. No statistical methods were used to initially determine sample size in these experiments. Experiments were not randomized, and investigators were not blinded to allocation during experiments and outcome assessments.

**Code availability**
Source code for all software and tools used in this study, with documentation, examples, and additional information, is available at the URLs listed above.

Fig. EV1 Esparza et al.

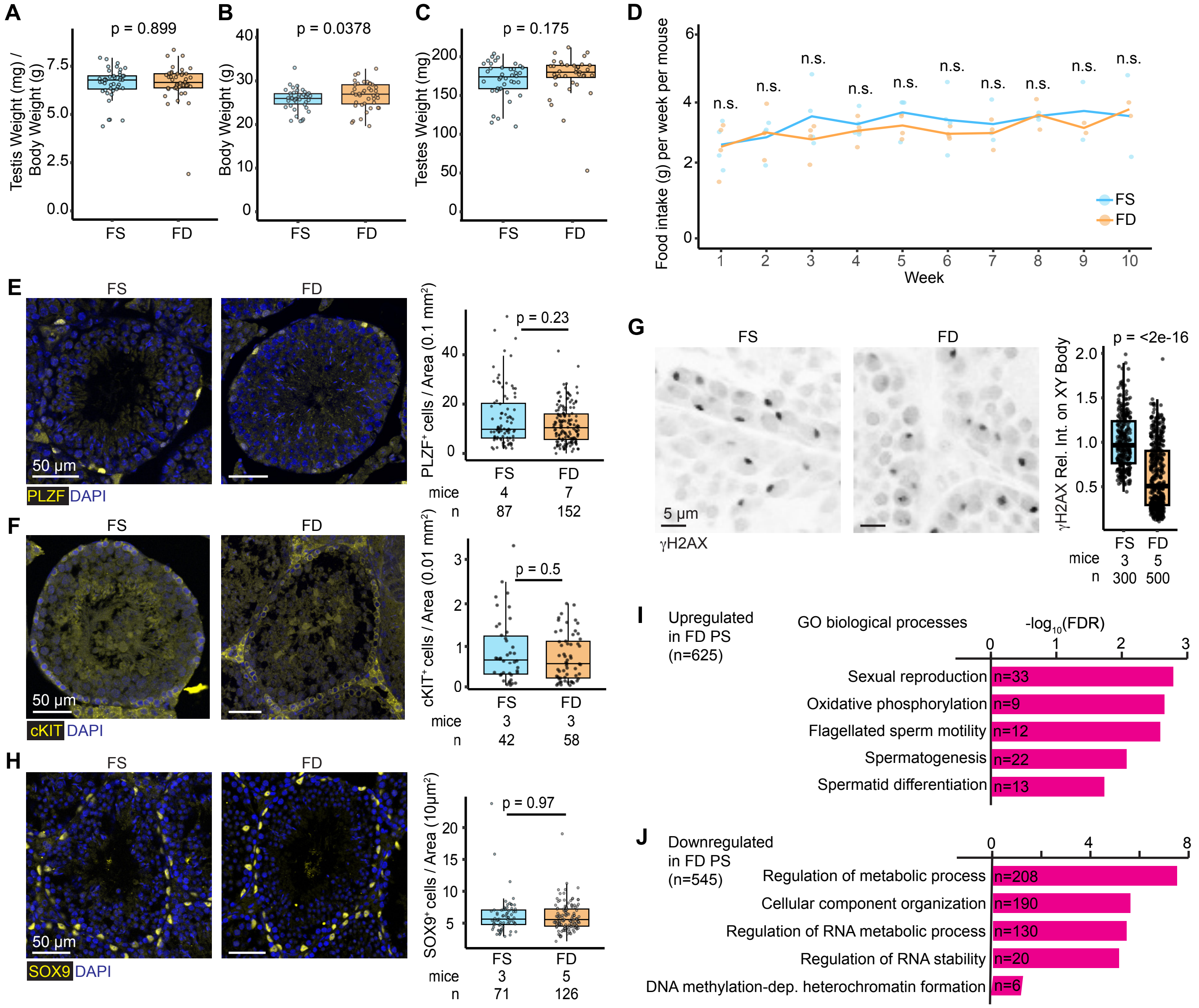

**Fig. EV1. Folate deficiency does not alter testis size or early germ cell populations but is associated with pachytene transcriptional changes.**

(A–C) Box-and-whisker plots comparing testes-to-body weight ratio, body weight, and testes weight between FS and FD mice (thirty biological replicates per condition).

(D) Weekly food consumption per mouse in FS and FD groups throughout the dietary exposure period. Points represent biological replicates (paired littermates), and lines indicate group means for each week. No significant differences in food consumption were detected between diets at any time point using the Benjamini–Hochberg method. adjusted for multiple testing.

(E) Immunostaining for ZBTB16 (PLZF) marking undifferentiated spermatogonia, with quantification (four biological replicates per condition).

(F) Immunostaining for KIT marking undifferentiated spermatogonia, with quantification (three biological replicates per condition).

(G) γH2AX immunostaining marking PSs, with quantification of XY body intensity normalized to DAPI (three biological replicates per condition).

(H) SOX9 immunostaining marking Sertoli cells, with quantification (three biological replicates per condition).

(I, J) Gene ontology analysis of biological processes enriched among upregulated (I) and downregulated (J) DEGs in FD PSs.

Fig. EV2 Esparza et al.

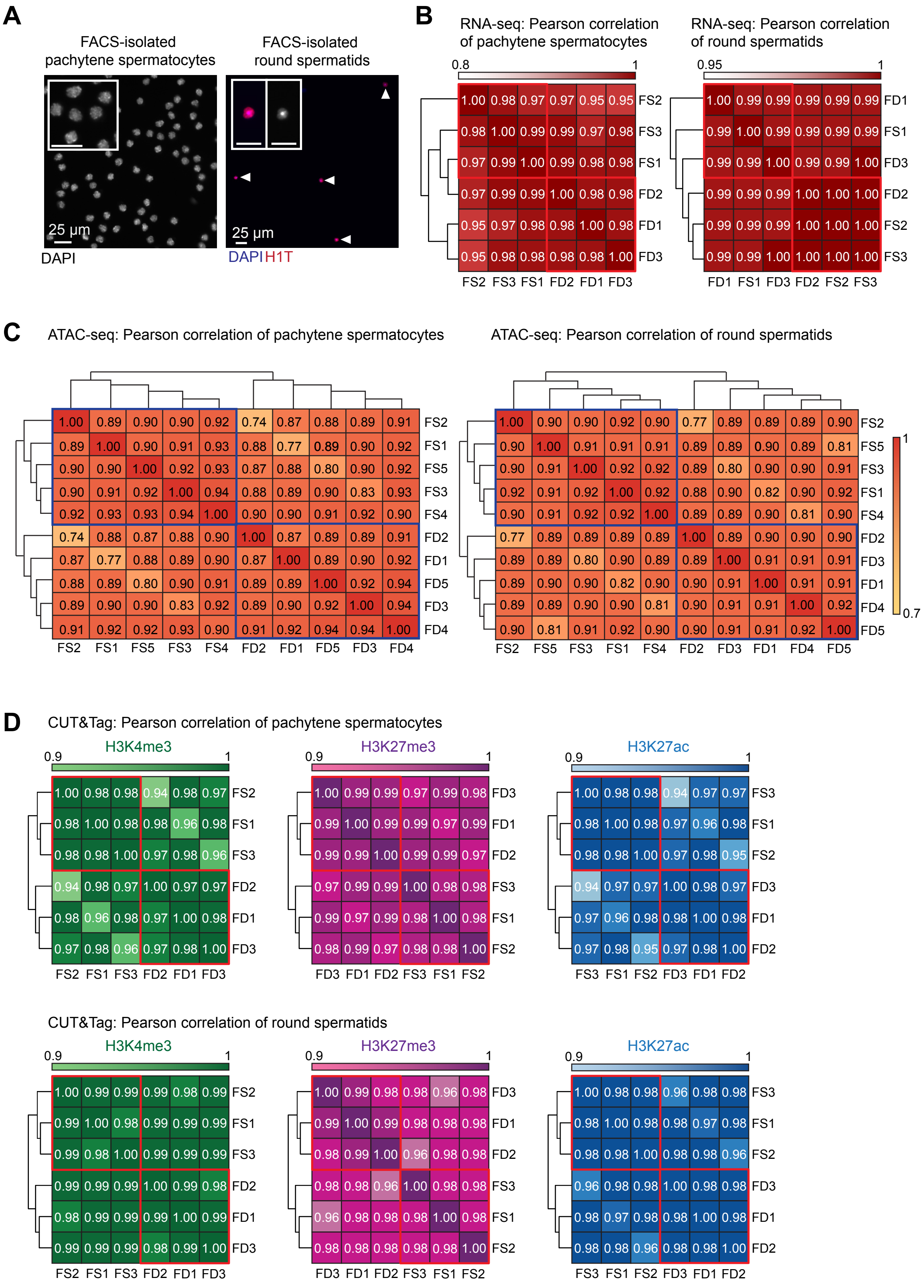

**Fig. EV2. Reproducibility of transcriptomic and epigenomic datasets.**
(A) Immunostaining of FACS-isolated PSs stained with DAPI (left) and RSs (right) stained with DAPI and H1T. Insets show higher magnification of PSs and RSs.
(B) Pearson correlation of RNA-seq datasets from FS and FD PSs (left) and RSs (right) (three biological replicates per condition).
(C) Pearson correlation of ATAC-seq datasets from FS and FD PSs (left) and RSs (right) (five biological replicates per condition).
(D) Pearson correlation of CUT&Tag datasets for H3K4me3, H3K27me3, and H3K27ac from FS and FD PSs (top) and RSs (bottom) (three biological replicates per condition).

Fig. EV3 Esparza et al.

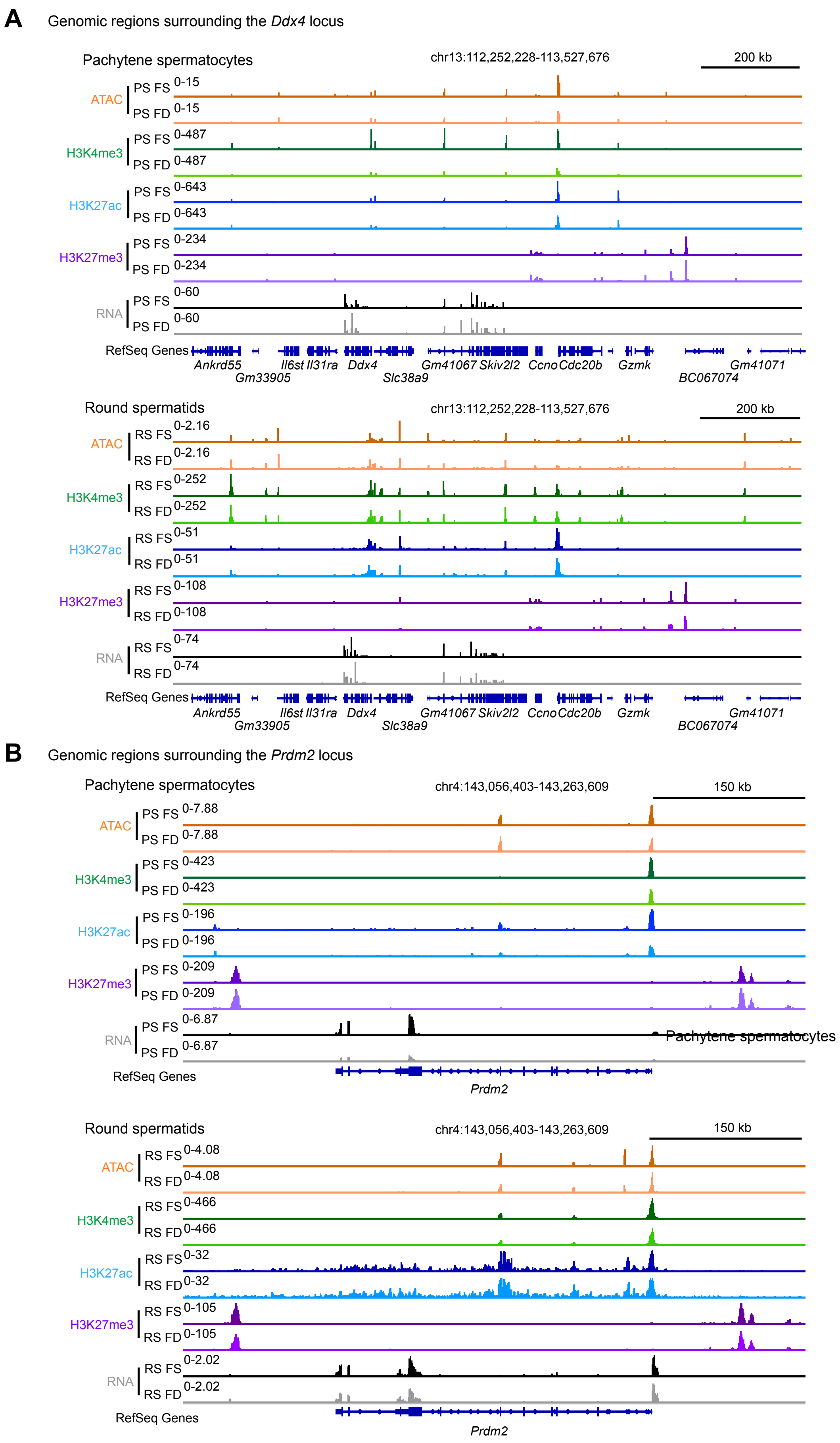

**Fig. EV3. Representative track views of ATAC-seq, CUT&Tag, and RNA-seq.**
(A) Genomic region surrounding the *Ddx4* locus.
(B) Genomic region surrounding the *Prdm2* locus.

Fig. EV4 Esparza et al.

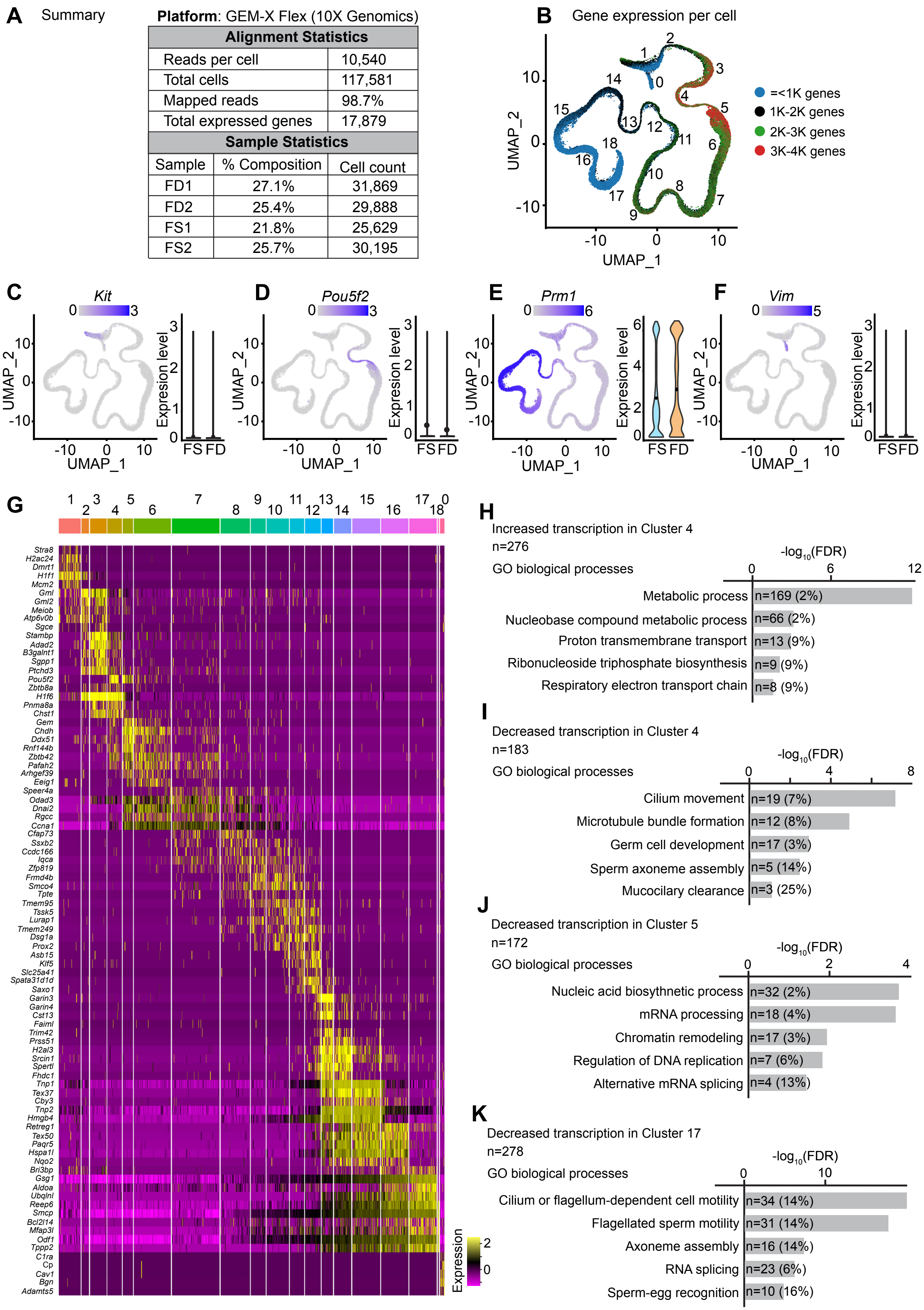


| Alignment Statistics | |
|---|---|
| Reads per cell | 10,540 |
| Total cells | 117,581 |
| Mapped reads | 98.7% |
| Total expressed genes | 17,879 |

| Sample | % Composition | Cell count |
|---|---|---|
| FD1 | 27.1% | 31,869 |
| FD2 | 25.4% | 29,888 |
| FS1 | 21.8% | 25,629 |
| FS2 | 25.7% | 30,195 |

**Fig. EV4. Quality control, cluster validation, and functional annotation of spermatogenic cell populations.**
(A) Summary of scRNA-seq quality control metrics.
(B) UMAP colored by number of detected genes per cell.
(C–F) UMAP feature plots of marker genes: *Kit*, *Pou5f2*, *Prm1*, and *Vim*.
(G) Heat map of the top five marker genes per cluster ordered by inferred developmental progression.
(H–K) Gene ontology analysis of DEGs in selected clusters.

Fig. EV5 Esparza et al.

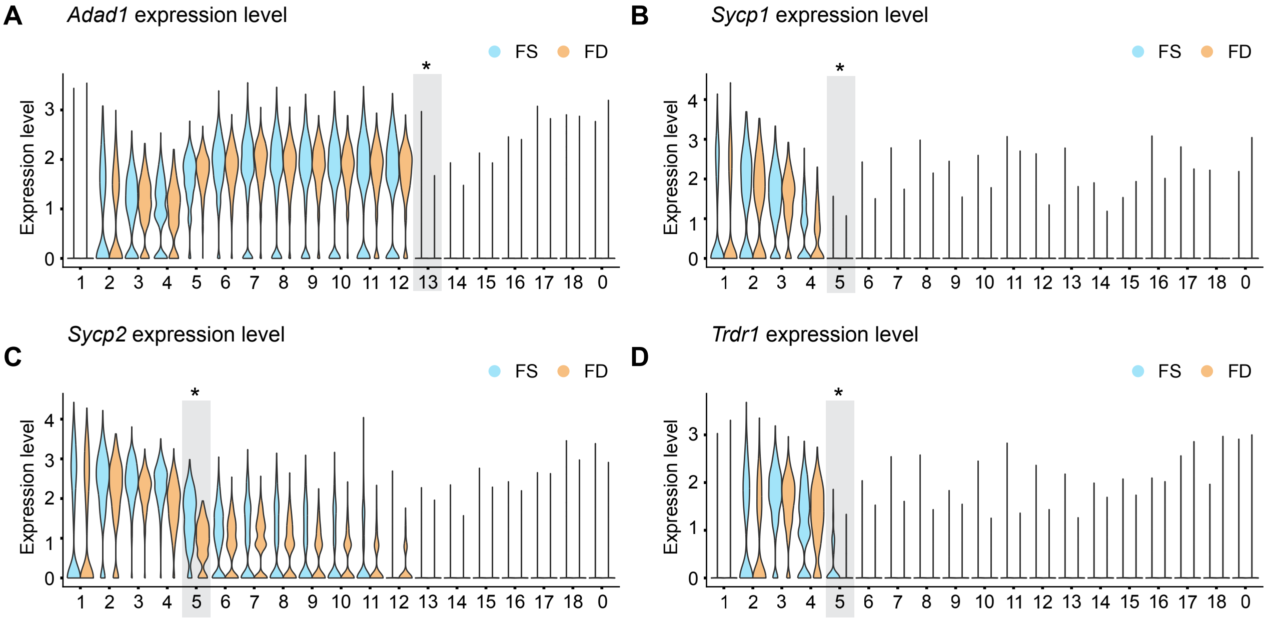

**Fig. EV5. Expression of germline reprogramming–responsive genes across spermatogenic cluster populations**
(A–D) Violin plots showing expression of *Adad1* (A), *Sycp1* (B), *Sycp2* (C), and *Tdrd1* (D) across spermatogenic cell clusters. Clusters highlighted in gray indicate those in which the gene was identified as a differentially expressed gene (DEG) in the scRNA-seq analysis.
